# From Tasks to Topology: Dorsal and Ventral Streams Emerge in Optimized Neural Networks

**DOI:** 10.1101/2025.11.16.688720

**Authors:** Tahsin Reza, Ewan Jordan, Steven T. S. Luo, Kirtan Patel, Jessica Tang, Matthias Niemeier

## Abstract

The primate visual system is organized into dorsal and ventral pathways, classically linked to visuomotor control and perception. A long-standing question is whether this division reflects intrinsic architectural priors or emerges from task demands. We trained a single convolutional network to perform classification and grasp prediction of 3D objects, without imposing modular structure. Dual-stream topology - functionally distinct visuomotor and perceptual pathways - emerged spontaneously with rich cross-communication. Shapley value analyses revealed that action- and perception-selective features developed progressively across depth, reflecting task-driven hierarchical specialization. Time-resolved EEG showed that model activity mapped onto dissociable temporal components in human cortex: ventral-aligned signals emerged early and late, where dorsal- and ventral-aligned responses coincided in the intervening interval. These results demonstrate that task optimization alone can explain core features of dorsal-ventral organization, and that distinct temporal roles for perception and action arise naturally atop a shared feedforward scaffold, without requiring architectural hard-coding or recurrence.

---

It is generally held that visual processing in the primate cerebral cortex is performed in two gradually diverging cortical streams, a dorsal stream projecting from striate and extra-striate areas to parietal cortex and a ventral stream extending from early visual layers into lateral occipital and inferior temporal regions (Figure 1A). The two streams have been associated with different functional distinctions, including spatial and object processing^1^, stimulus properties such as motion and form^2^, or relatively more egocentric versus allocentric coding^3,4^. Others have challenged the idea of a strict separation^5–10^, consistent with evidence for interactions between dorsal and ventral areas at multiple stages^11–13^. Still others have argued that each stream is itself further separated into substreams^14,15^. More recently, it has been suggested that visual streams arise from topographic principles and self-supervised learning that extracts statistical regularities embedded in the visual input, yielding data-driven multi-purpose representations^16^.

**Figure 1.**
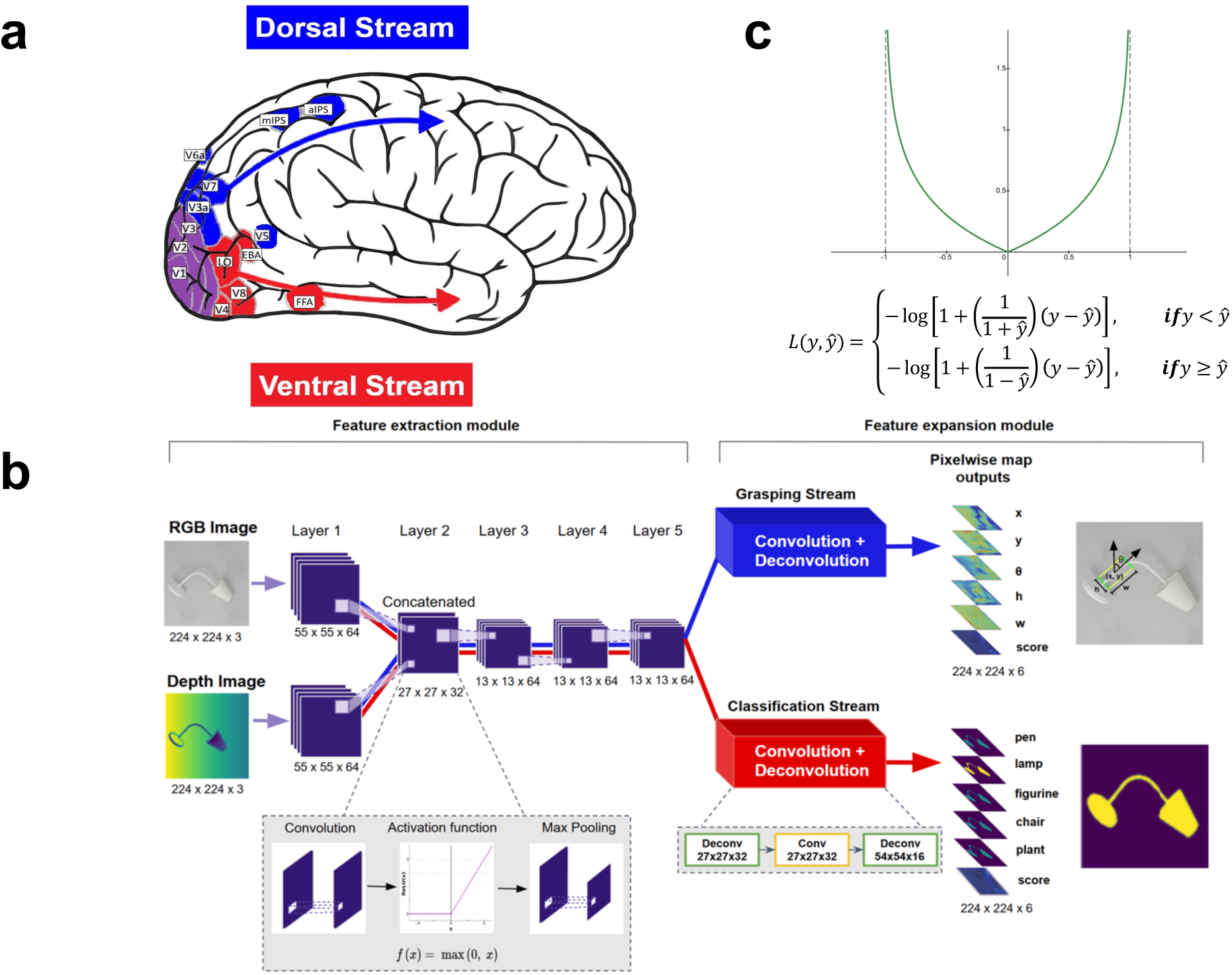
Dual-stream architecture in biological and artificial systems. **a.** Dorsal and ventral stream as hypothesized for the primate (here: human) cerebral cortex. **b.** Architecture of ANN used in this study. The network was jointly trained on a visuomotor task (grasping) and a perceptual task (object classification). Training was based on images (224 by 224 pixels) of real-world objects drawn from five different object categories^32^. Images included three color channels (red, green and blue, RGB) and a depth (D) channel, important for grasping^33^. The RGB and D channels are shown separately (left). The model was designed to capture depth-based spatial representations, emphasizing stereoscopic cues known to engage dorsal stream areas via V6A, a major source of visuospatial input to the dorsal system^34^. We deliberately omitted motion information^2^ to reduce computational complexity and to isolate the contribution of depth structure to dorsal processing. Depth and color channels were provided as separate inputs to a convolutional Feature extraction module with 5 layers, similar to other convolutional neural networks (CNNs)^35^. In this variant of CNNs, the color and depth channels were processed independently in the first layer and merged in the second to form joint representations integrating color and depth information. Each layer of the network applied a set of spatial filters to produce retinotopically organized feature maps, where each feature map corresponded to a population of units with identical tuning (e.g., all preferring oblique edges, see feature map 10, Fig. 2A) but with receptive fields at different spatial positions. To mimic the nonlinear response properties of neurons, each feature map was followed by a rectified linear (ReLU) activation, and in layers 1 and 2, by a max-pooling operation that reduced spatial resolution and increased receptive field size in the subsequent layer. The RGB portion in Layer 1 was pretrained on ImageNet^36^, and has been shown to resemble response properties of area V1^35^. Crucially, the Feature extraction module formed the shared backbone of the network that was used by both grasping and classification computations. It was not explicitly segregated by design; instead, dorsal- and ventral-like pathways emerged purely through task-based optimization. Two expansion modules, each consisting of convolutional and deconvolutional layers, were used to efficiently train the ANN on grasping and object classification. These modules served as computational constructs rather than a biological mechanism. For grasping, the expansion module provided an efficient means of mapping a discrete set of labelled grasp examples onto the continuous space of correct grasp configurations^31^. Specifically, the output of the Grasp module consisted of five “heatmaps” with the same dimensions as the input images, encoding parameters of bounding boxes that defined individual grasps: x, y, θ, h, and w, corresponding to horizontal and vertical position, orientation, height, and width of the grasp, respectively. An example bounding box is shown on the right. These parameters could be submitted to a motor control module to generate grasp movements in a biological or robotic effector^31^. They represent the minimal set of variables that any grasping, biological or artificial, must compute explicitly or implicitly^37^. A sixth score map quantified the predicted quality of the individual grasps. The Classification module had an architecture matched to the Grasping module to ensure comparable computational complexity. It also produced five output maps, each corresponding to one of the object categories used in training plus a score map. I.e., for each input image the ANN was trained to activate the map associated with the object class such that only pixels overlapping the object were active. Methods for details. **c.** A custom double-log loss function was used to jointly optimize both tasks. This novel loss function combines properties of mean squared error and cross-entropy loss, making it suitable for simultaneously training continuous (grasp) and categorical (classification) outputs. See Methods for full specification.

Among this diversity of views, the framework proposed by Goodale and Milner^17^ stands out because it ties the division of visual streams to ecologically relevant goals. It posits that visual representations are fundamentally task-driven - the ventral stream is optimized for visual perception, whereas the dorsal stream is optimized for the visual control of action. Thus, from an evolutionary perspective, it is easy to imagine selective pressures favoring neural systems that allow an animal to recognize what something is and to act upon it effectively. By contrast, it is less clear how selection would favor a separation based on purely abstract stimulus categories such as “objects” or “space,” which do not themselves constitute goals.

The perception and action model’s two task-based optimization principles imply a clear prediction: if these principles are used as teaching signals in a neural network, biological or artificial, the network should spontaneously form distinct processing streams for action and perception. It is striking that, to date, there have been only limited attempts to test this prediction empirically. Two studies have reported dorsal- and ventral-stream-like properties emerging in simulated networks, but in both cases the dual-stream architecture was hard-wired and trained to represent stimulus properties^18,19^ (for a model with triple-stream architecture see^16^). Ecologically relevant, task-based optimization was instead used in a study^20^ that trained a network to predict self-motion required for the ability to orient oneself, and important property of the dorsal stream^21^. However, that study did not employ a task-related optimization principle associated with the ventral stream. Other studies used task-based optimization for both visual streams but again within networks containing pre-designed processing branches^22–25^. Such modular organization with functional specialization has been shown to emerge in ventral substreams through task optimization even without architectural constraints^26^. By contrast, others have argued that data-driven self-organization alone is sufficient to explain large-scale visual-stream structure^16^. However, that view effectively treats visual cortex as impervious to task-driven optimization. Yet the brain - including visual cortex - is necessarily plastic to environmental and behavioral pressures. If task demands influenced representational tuning, then optimizing a single network for distinct perceptual and visuomotor goals should itself induce segregation.

To test this central prediction of the perception-and-action framework proposed by Goodale and Milner^17^, we trained an artificial neural network on two distinct visual tasks, one for object classification and one for action, without imposing any architectural modularity. We show that dual-task optimization alone is sufficient for a functional division to emerge, even without topographic constraints, yielding an integrated organization with both segregation and cross-communication reminiscent of dorsal- and ventral-stream processing. Moreover, recording high-density electroencephalography (EEG) in human participants, revealed that activity in dorsal and ventral streams diverges not only across brain regions but also in temporal dynamics, suggesting that time constitutes a key axis of their functional organization. Together, these findings support the idea that differences in task demands, rather than predefined anatomical or statistical constraints, may drive the functional organization of visual cortex. In this view, the visual cortex may first acquire broadly multi-purpose representations, which then differentiate under the optimization pressures imposed by perceptual and action goals.

## Results

### From shared to specialized computations

We trained a custom-made dual-task convolutional neural network (Fig. 1b; Methods) to perform two tasks: predicting grasp points on objects and classifying the objects. These tasks are associated with more dorsal or ventral brain regions, respectively^27–30^, with grasping relying on three-dimensional structure derived from stereoscopic depth cues^34,35^. To isolate this depth-based component of dorsal processing, the network received color and depth input. Motion information was omitted to reduce computational complexity. The network comprised one Feature extraction module that was jointly optimized using two task-specific expansion modules, one for grasping and one for object classification, and a custom double-log loss function (Fig. 1c) that balanced continuous and categorical learning objectives. The grasping module used a convolutional-deconvolutional expansion head, following established designs for pixel-wise prediction of grasp configurations^31^, whereas the classification module used an analogous expansion head rather than conventional fully connected layers. This design ensured comparable computational complexity between the two tasks and preserved spatial structure in both outputs.

To examine how the Feature extraction module of the network internally represented the two tasks, we quantified the marginal contribution of each feature map in each of its five layers to grasping and classification performance using Neuron Shapley values^38^ (e.g., Fig. 2a). Correlations between these task contributions were significant in the first two layers (r ≥ 0.65), indicating shared low-level computations. However, the correlations dropped sharply from layer 2 to 3 and remained low through layers 3-5 (Fig. 2B). Importantly, correlations were near zero across deeper layers, with layer 5 showing a numerically negative but nonsignificant correlation. Together, these results, from a shared Feature extraction module without built-in task segregation, suggest that task-related specialization unfolded through orthogonalization rather than antagonistic coding.

**Figure 2.**
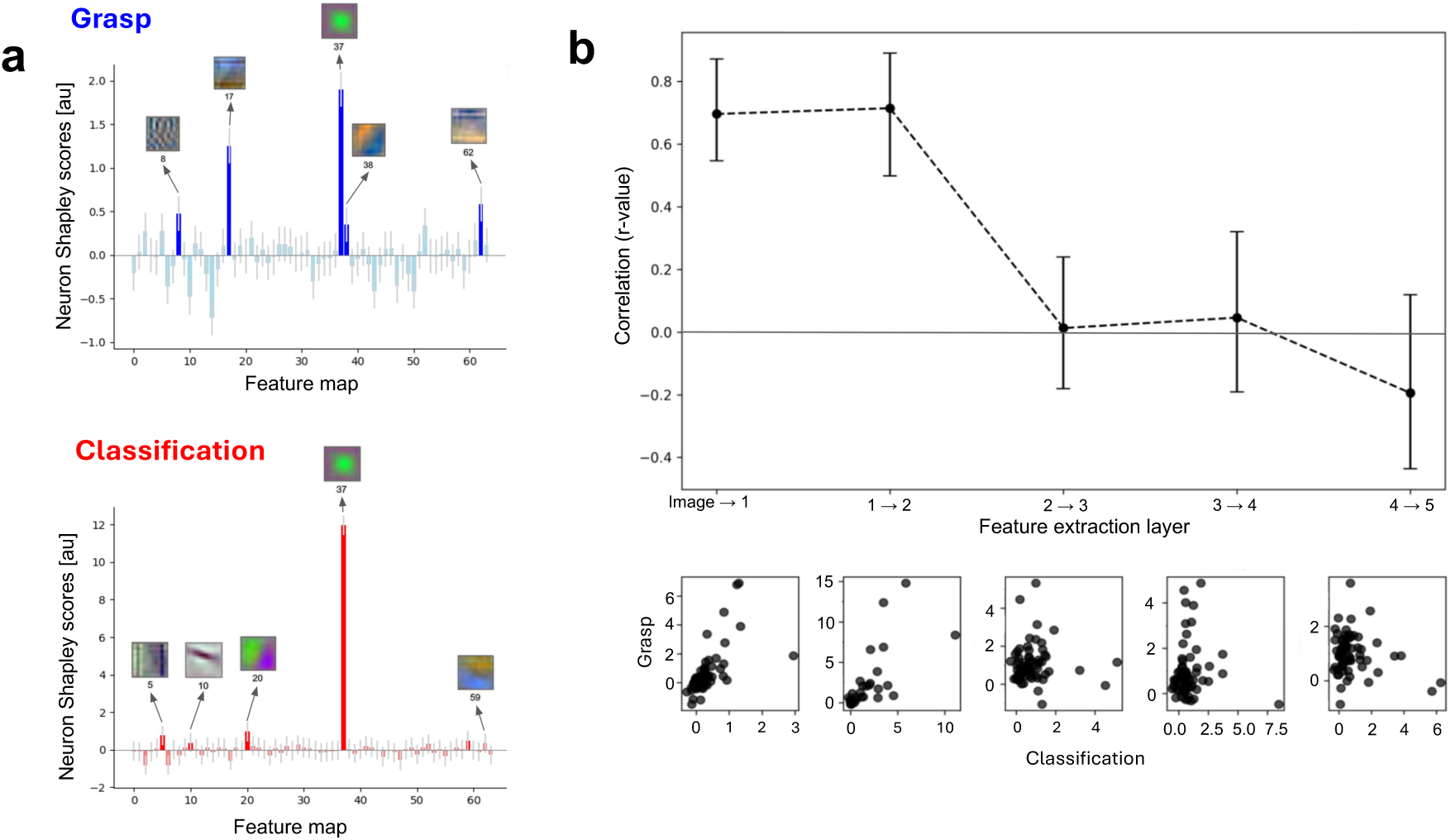
Task-selective contributions of feature maps quantified using Neuron Shapley scores. **a.** Neuron Shapley scores for individual RGB feature maps in Layer 1 of the shared Feature extraction module. Scores reflect each map’s unique contribution to task performance, computed by iteratively measuring performance drops following the removal of different combinations of feature maps. Higher scores indicate stronger task relevance. Insets show preferred input stimuli, derived by activation maximization, for the five feature maps most strongly contributing to each task’s performance. **b.** Top: correlations between Neuron Shapley scores for grasping and classification across all five shared layers of the Extraction module. Error bars: bootstrapped 95% confidence intervals. Bottom: scatter plots of Neuron Shapley scores for grasping and classification in each of the 5 layers.

The stepwise decorrelation further indicates that a distinct representational bifurcation emerged spontaneously within the Feature extraction module of the network, driven solely by the differing demands of the two tasks. The abrupt transition from shared to independent feature use is consistent with separate processing pathways for action and perception arising in the brain. At the same time, the lack of significant negative correlations in deeper layers shows that this division does not reflect strict anatomical segregation, but rather allows for cross-talk, consistent with the interplay observed between the biological streams.

### Double dissociations

A defining hallmark of dorsal-ventral stream separation comes from lesion studies: damage to ventral areas impairs object recognition while sparing reach and grasp movements^28,39,40^, whereas dorsal lesions yield the opposite pattern^41–43^. Such lesion findings, unlike correlational evidence, reveal the causal contributions of each pathway to action and perception. In biological brains, however, lesions can lead to a combination of direct deficits and compensatory processes that obscure the underlying mechanisms^44^. By contrast, such adaptive nonlinearities are absent in our network, offering a more transparent demonstration of how dissociable functional streams can arise from task optimization alone. To test whether a comparable double dissociation would emerge, we identified feature maps according to their causal contribution to each task, as quantified by Neuron Shapley values, and progressively removed those with the largest grasp- or classification-specific impact. This data-driven lesioning procedure identifies task-critical units based on function alone without assuming predefined modules. A double dissociation was operationally defined as a lesion type producing a significant impairment in one task but not the other, and the opposite pattern for the complementary lesion type.

For the first layer, we found little evidence for a double dissociation (Fig. 3a). Removing more than one grasp-impact map (≥0.7% of layer 1) significantly disrupted grasp performance, while classification remained intact even after 50% damage. However, network performance was nearly equally sensitive to classification-impact lesions, which impaired classification and grasping after only 0.7% and 1.4% damage, respectively. Similarly, even small classification-impact lesions (3.1-6.2%) to layer 2 significantly impaired both tasks (Fig. 3b), whereas a narrow range of grasp-impact lesions (12.5-17.5%) selectively disrupted grasping. Thus, damage to early visual processes was unlikely to produce clear double dissociations.

**Figure 3.**
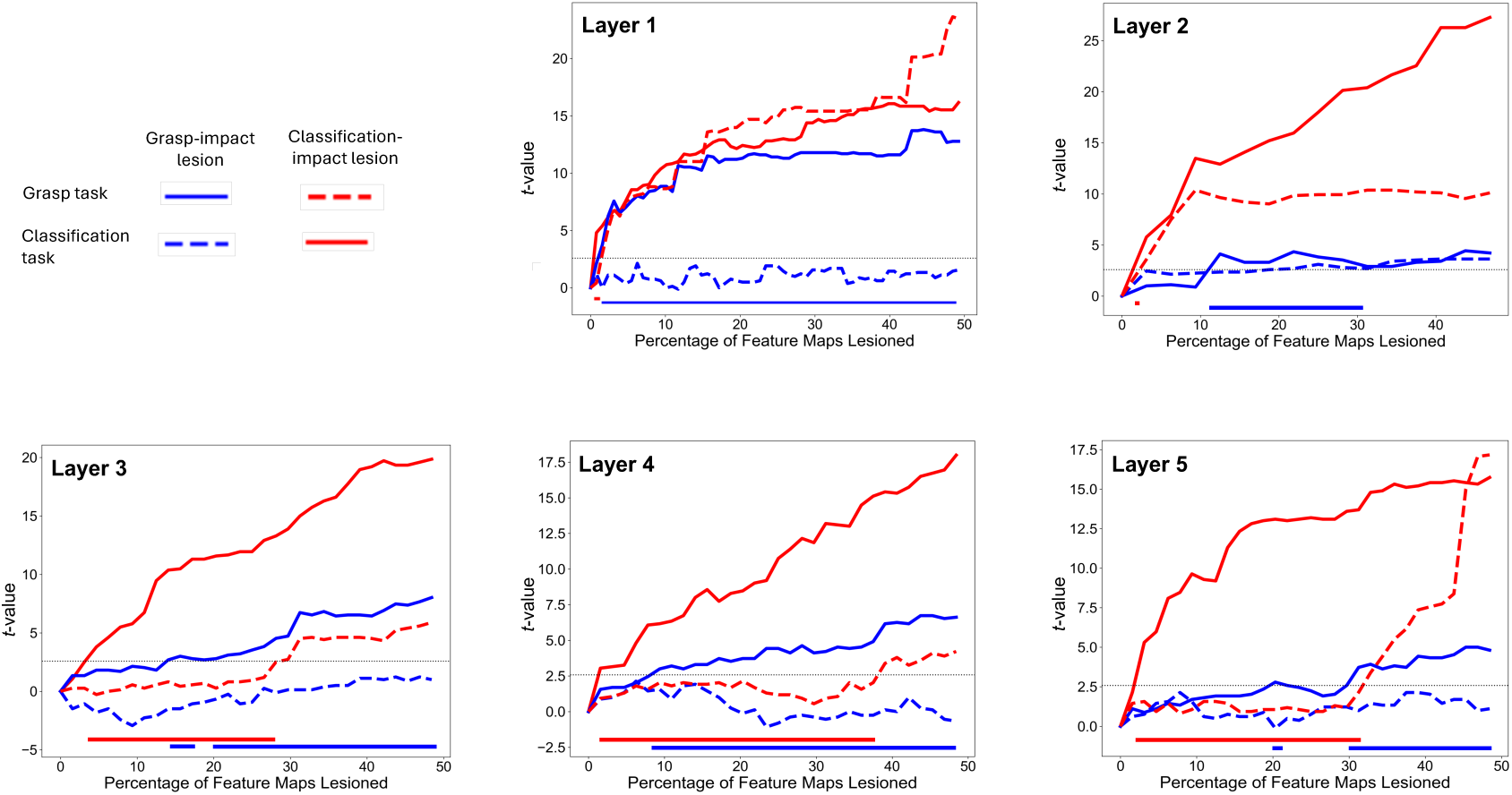
Double dissociations. Task performance as a function of progressively lesioned networks. Red and blue curves show t-values for normalized grasping and classification performance, respectively, relative to intact performance. Lesions targeted features with greater Neuron Shapley scores (i.e., that were more important) for grasping than classification, or vice versa. Black dashed line marks level of significance (p < 0.05). Blue and red horizontal bars mark lesion ranges where the grasp task but not the classification task is significantly impacted or vice versa.

By contrast, marked double dissociations emerged in the deeper layers (Fig. 3c,d). In layer 3, lesions affecting ≥14% of grasp-impact maps impaired grasping but not classification, whereas classification-impact lesions between 4% and 27% selectively reduced classification accuracy. In layer 4, grasp- and classification-impact lesions exceeding 9% and 2%, respectively, produced clear task-specific deficits. A similar pattern was observed in layer 5, where lesions of ≥30% grasp-impact units impaired grasping alone, while 3-33% classification-impact lesions disrupted classification alone.

Together, these findings demonstrate that a double dissociation between perception and action computations emerged spontaneously in the network’s deeper representations, mirroring the functional segregation of dorsal and ventral streams in the lesioned brain. This convergence of representational divergence and causal dissociation justifies referring to these pathways as dorsal- and ventral-like in subsequent analyses, including their cross-stream connectivity.

### Inter-layer connectivity

So far, we examined whether computations within each layer shared resources for grasping and object classification or whether these resources were separate. This leaves open how information flows between layers: whether features that support action preferentially project to action-relevant features at the next level, and likewise for perception, or whether connectivity remains broadly shared. Dual-stream accounts and hierarchical vision models both imply that such specialization, if present, should strengthen with depth, as early stages extract general visual structure and later stages acquire task-specific features. We therefore tested whether inter-layer connections show enrichment for within-task routing and whether any such bias increases across layers.

To address this, we next asked whether connections between layers preferentially linked feature maps with similar or different task specializations. Whereas the lesion analysis ranked maps by their functional impact on each task, connectivity analysis requires identifying feature maps with task selectivity. We therefore categorized maps by task bias, labeling them as dorsal-biased or ventral-biased when their Neuron Shapley contributions to grasping versus classification differed by a standardized effect size. A stricter cutoff (|d| ≥ 0.5) isolated strongly biased maps, while a more inclusive threshold (|d| ≥ 0.15) captured weaker preferences. Maps with smaller effect sizes were considered non-selective. Inter-layer connections between feature maps were then categorized as (i) dorsal-dorsal, (ii) ventral-ventral, (iii) cross-connections (dorsal→ventral or ventral→dorsal), or (iv) unspecific (involving at least one non-selective map).

As shown in Figure 4a, distinct dorsal and ventral connectivity patterns emerged, especially in deeper layers. The corresponding pie charts (Fig. 4b, top row) illustrate how the proportional distribution of connections shifted with depth; within-stream (dorsal-dorsal and ventral-ventral) connectivity increased alongside a concurrent rise in cross-connections, whereas unspecific connections became less frequent. Consistently, weighted χ² tests revealed significant connectivity gradients for within-stream connections (dorsal and ventral) at deeper transitions, specifically from layer 2→3 relative to 3→4, and from 3→4 relative to 4→5. Furthermore, cross-connections became significantly stronger from 2→3 relative to 3→4 and remained comparably strong thereafter. We obtained similar results when using a less conservative effect size threshold (d ≥ 0.15, Fig. 4b, bottom row) or a non-parametric rank-based criterion (top k; Fig. 4c).

**Figure 4.**
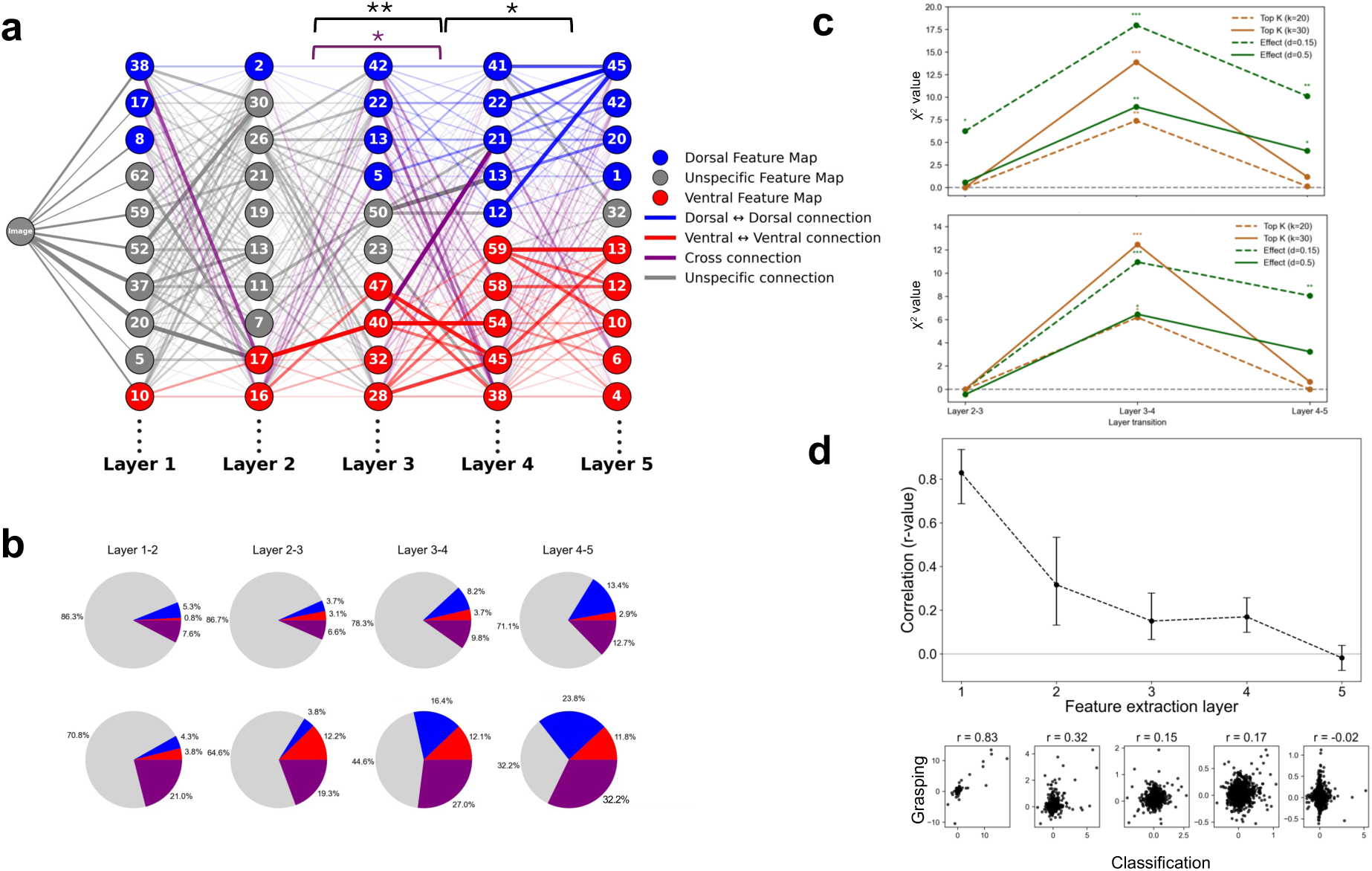
Inter-layer connectivity. **a.** Functional connectivity diagram illustrating information flow between the five feature-extraction layers of the ANN. For visibility, nodes and edges were thresholded. Nodes represent feature maps with the top ten Neuron Shapley scores for grasping or object classification. Feature maps were categorized by task-specificity based on effect size: d = |Shap(grasp) - Shap(cls)|/|Shap(grasp) + Shap(cls)|, with d ≥ 0.5. Edges represent weighted directed connections, color-coded by task alignment (dorsal-dorsal, ventral-ventral, cross-connections, or unspecific). Edge opacity scales with the absolute connection weight between feature maps. Brackets above the graph indicate layer transitions with significant connectivity gradients between successive layers (e.g., layer 1→2 vs. 2→3; χ² test, p < 0.05). (Connections from input→layer 1 were classified as unspecific and therefore excluded from the gradient analyses in B-C.) Black brackets mark gradients where within-stream (dorsal + ventral) connectivity increased relative to cross + unspecific connectivity; the purple bracket marks a significant gradient for cross-connections relative to unspecific connections. **b.** Pie charts summarizing the distribution of connection types between successive layers, weighted by the average Neuron Shapley scores of the connected feature maps. Top row: conservative criterion for node categorization (d ≥ 0.5). Bottom row: liberal categorization criterion (d ≥ 0.15). **c.** Summary of χ² results testing for significant connectivity gradients. Top graph: χ² statistics for within-stream connectivity gradients based on the conservative and liberal effect-size criteria as in A. and B. as well as an additional rank-based (top-k) criterion (Methods). **d.** Approximate connection-level Shapley values based on SVARM^45^, plotting the correlation between grasp- and classification-related connectivity for each layer pair. Top graph: correlations between grasp- and classification-related connection importance as a function of all layer transitions, including input→1. Bottom insets: scatter plots of Shapley scores for grasp-versus classification-related connections within each layer pair.

To provide a complementary perspective, we next estimated connection importance directly, using an approximate Neuron Shapley method applied to the weights themselves. Because of the large number of connections, the approach relied on a computationally efficient estimation rather than an exact Shapley calculation. Nevertheless, this analysis provided converging evidence for gradual separation of dorsal- and ventral-like connectivity. Specifically, connections from the input to layer 1 showed highly similar importance for both tasks (r = 0.83), with correlations declining across deeper layers (Fig. 4d).

In sum, these inter-layer analyses parallel the within-layer findings, revealing a progressive differentiation of dorsal- and ventral-like streams with network depth. At the same time, the persistence of cross-connections underscores that this functional division is not achieved through strict segregation, but through coordinated specialization with continued interaction between streams.

### ANN vs. human brain comparisons

Having established that the ANN exhibited a functional segregation reminiscent of the biological dorsal and ventral streams, we next compared simulated network activity with human brain responses. To this end, we conducted an electrophysiological experiment, recording high-density EEG while participants performed grasping and classification tasks with physical three-dimensional objects (Fig. 5a). Event-related potentials (ERPs; Fig. 5b) were used to compute representational dissimilarity matrices (RDMs) across consecutive 25-ms time windows with equivalent RDMs derived from activations in the five layers of the ANN’s Feature extraction module (Fig. 5c). Comparing ERP RDMs and model RDMs produced significant correlations for both tasks (Fig. 5d). These correlations were substantial (up to r ≈ 0.3), given that they reflected raw values that were not normalized to a noise ceiling, suggesting that the task-optimized ANN converged on representational geometries characteristic of the human brain.

**Figure 5.**
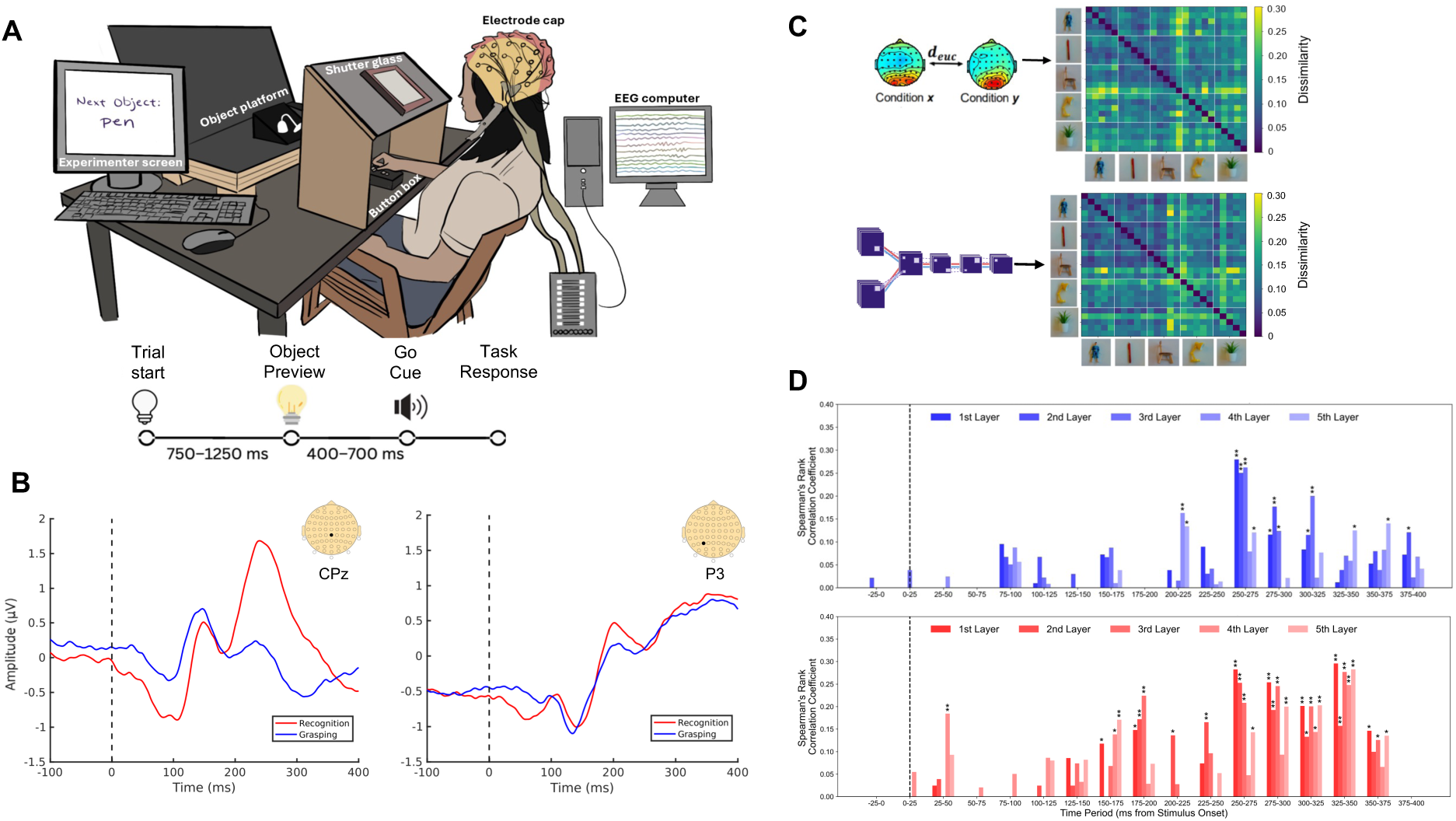
Comparing ANN activation with electrophysiological responses in the human brain. **a.** Experimental set-up of the EEG experiment (n = 16 participants, 8-9 hours per participant). Participants viewed three-dimensional objects, one at a time, from the five object categories for 400 to 700 ms. Following an auditory cue, they picked up the object with a precision grasp or classified it verbally. **b.** Grand-averaged ERPs recorded over human cortex. **c.** Top: Example representational dissimilarity matrix (RDM) derived from ERPs by calculating Euclidian distances between postero-central voltage patterns across all experimental conditions, computed in 25 ms time windows. Bottom: Corresponding RDM calculated from the ANN’s Feature extraction module, based on activations from layer 1 to RGB-D images digitized from the same objects used in the human experiment. **d.** Spearman rank correlations between EEG- and model-derived RDMs as a function of preview time. Each bar represents the correlation between ERP RDMs within specific time windows and model RDMs from the five Feature extraction layers of the ANN.

Strikingly, the temporal dynamics of these correlations differed systematically between tasks. For grasping, earliest significant correlations arose at 200-225 ms, peaking at 250-275 ms and gradually declining thereafter. In contrast, object-classification correlations emerged earlier (150-175 ms), peaking at 175-200 ms, and persisted over an extended period from 225 to 375 ms, with additional peaks at 250-275 and 325-350 ms.

These peaks map onto established electrophysiological components. The earliest classification-related peak (175-200) coincides with the N2 ERP component, previously linked to recurrent mid-level feature refinement^46^ and early category discrimination in ventral visual areas^47^. The subsequent joint peak for both tasks (250-275 ms) coincides with the P3 complex, associated with cognitive updating and a feedback pass through higher-order visual and parietal regions^48^, consistent with reactivation of task-relevant object features^49,50^. Finally, the later classification-specific peak (300-325 ms) likely indexes early access to semantic object representations^51–53^. Together, these latencies point to recurrent or feedback mechanisms dominating representational similarity between model and brain activity.

Notably, the ANN itself is a strictly feedforward architecture, lacking any explicit recurrence. Yet, the timing of the correlations mirrors a cascade from early to higher-order stages: the first two peaks were more strongly associated with activity in the ANN’s early-to-mid layers (layers 1-3), whereas the late classification-specific peak showed a more uniform distribution of correlations across all feature-extraction layers. This correspondence suggests that hierarchical, feedforward computations in the model recapitulate the temporal signatures of recurrent processing observed in the brain.

## Discussion

Our results demonstrate that functional segregation between action and perception can emerge spontaneously from task optimization alone, without any imposed architectural priors^18,19,22–25^. The resulting dual-stream organization reproduced hallmark properties of the biological dorsal and ventral pathways, specialized computations emerging hierarchically over successive processing stages and unfolding over distinct temporal scales.

This finding provides direct computational support for the framework proposed by Goodale and Milner^17^, which attributes the division of visual processing to task-specific optimization principles rather than anatomical separation per se. In our simulations, the network’s dual-task training gave rise to dorsal- and ventral-like streams with characteristic specialization for visuomotor and perceptual processing, respectively. Crucially, these specializations coexisted with extensive cross-connections, demonstrating that functional segregation and integration are not mutually exclusive but emerge jointly from shared computational constraints.

In this way, the model reconciles seemingly opposing perspectives in the literature, those emphasizing dorsal-ventral segregation^1,17^ and those emphasizing cross-stream interactions^5,8,11–13^, by showing that what have been framed as opposing accounts instead reflect complementary consequences of the same ecological optimization pressures.

According to this interpretation, action optimization naturally entails spatial coding^1^, as accurate grasp planning requires representing object position and geometry in egocentric coordinates. On the other hand, object classification is supported by view-invariant representations that approximate allocentric coding^4^. While our model did not explicitly measure coordinate frames, the emergent division of labor is consistent with an interpretation of the dorsal-like pathway supporting behavior in body-centered space and the ventral-like pathway supporting object identity abstraction. Thus, optimization for distinct behavioral goals may naturally give rise to the representational reference-frame distinction hypothesized in dual-stream theories, connecting egocentric action guidance with allocentric object identification.

In parallel, recent modeling work has argued that large-scale visual-stream architecture is sufficiently explained by self-supervised learning on natural image statistics under topographic constraints, without reliance on task-specific objectives^16^. In contrast, our results show that task-driven visual-stream segregation is inevitable, given that the visual system is not impervious to environmental and behavioral pressures. This framework challenges input-driven explanations by emphasizing that a system capable of extracting latent visual structure presupposes adaptation to those same pressures. In this view, topography and statistical learning are not alternative mechanisms but emergent consequences of task-driven optimization.

A notable feature of our findings is that the temporal segregation between visuomotor- and perception-aligned signals in human EEG emerged even though our network architecture was purely feedforward, lacking recurrence or temporal dynamics. Prior research has reported that early layers in deep networks tend to correspond to earlier EEG or MEG responses while deeper layers corresponded to later responses^54–57^, traditionally viewed as evidence for feedforward hierarchies unfolding over time. We observed a modest manifestation of this pattern, with earlier and intermediate peaks of model-brain correspondence showing stronger correlations with early-to-mid network layers. Importantly however, our earliest correspondence peak still fell within timeframes typically characterized by recurrent and feedback-dominated visual cortex activity. Thus, correlations between network depth and neural timing do not imply that the brain is operating in a strictly feedforward fashion. Instead, the fact that task-specific temporal structure emerged in a feedforward-only model suggests that optimization-driven hierarchical refinement can act as a scaffold that biological recurrence subsequently expands and stabilizes. In this view, recurrence may amplify, refine, and propagate distinctions already latent in feedforward processing, rather than generating them from scratch.

Beyond this feedforward scaffold, the EEG-ANN comparisons revealed a second dimension of organization. Not only did a dorsal-ventral differentiation emerge as a matter of computational representation, but also as different temporal dynamics for action and perception. Ventral-aligned signals rose earliest and re-emerged later, whereas dorsal- and ventral-aligned responses coincided during the intervening period, pointing to a temporal specialization that was not explicitly programmed into the model. This pattern is consistent with the idea that biological recurrence imposes a temporal ordering over pathways that are architecturally intertwined, such that perceptual evidence is rapidly extracted, visuomotor transformations are engaged as soon as sufficient structure is available, and higher-order perceptual refinements return later.

Time-resolved electrophysiology studies now show that dorsal and ventral pathways follow different temporal trajectories, with stream-specific signals emerging at distinct moments during visual processing. Some findings indicate earlier dorsal responses, others show early ventral selectivity^58–61^. Regardless of these differences between studies, there is agreement that dorsal and ventral processing follow partially dissociable schedules.

In this light, our observation of an early ventral-aligned peak, followed by a period of coinciding dorsal and ventral alignment and a later ventral peak, provides a concrete example of task-dependent temporal specialization. Rather than reflecting a fixed order of processing, these dynamics point to a flexible pattern of separation and synchronization as a central principle of dual-stream organization. Taken together, these findings suggest that the two streams are differentiated not only in function but also in the temporal roles they adopt as visual computations unfold.

Beyond temporal dynamics, another key property of dual-stream organization concerns its topographic expression in cerebral cortex. Here we use *topology* to refer to the network’s functional connectivity structure, distinct from *topography*, the spatial layout of cortical tissue. In the biological brain, functional specialization interacts with spatial proximity constraints that enable rapid and metabolically efficient communication^62^. Our model, by design, lacked such topographic constraints. Nevertheless, it developed distinct visuomotor and perceptual feature clusters. This suggests that cortical topography need not precede specialization; rather topography may co-develop with functional differentiation under shared pressures for efficient wiring and coordinated processing. Put differently, the brain’s spatial organization does not reflect the segregation of pre-formed modules but the manifestation of distributed functional gradients within a physically constrained substrate.

While our model captures central computational principles of dual-stream organization, it remains necessarily simplified in its sensory inputs. Biological vision integrates multiple dynamic cues, including motion and binocular disparity, to support action control in three dimensions. Our model incorporated stereoscopic depth information (RGB-D), consistent with the fact that grasping is fundamentally a 3D problem. Human psychophysics and robotics both demonstrate that removing binocular vision or relying solely on 2D inputs markedly impairs grasp performance. Indeed, grasping without depth cues yields measurably reduced grasp performance^34^, and contemporary robotics systems routinely incorporate RGB-D sensing for robust autonomous grasping^31,38^. Furthermore, neurophysiological research shows that stereovision serves as a major input to posterior parietal cortex^34^.

Motion signals, on the other hand, were not included in the present study. This was a pragmatic decision to isolate the core computational question, whether dual-stream organization emerges from task optimization, without compounding architectural and training complexity. Classic neurophysiology shows that motion provides an important input to dorsal stream computations^2^ and, several modelling efforts have demonstrated emergent dorsal-like specialization from dynamic, albeit monocular, visual input^18,19^. Our results complement these approaches by isolating a different biological constraint: the requirement to guide action in three-dimensional space. Importantly, motion is not synonymous with the dorsal stream; rather, motion is one of several privileged cues that the dorsal system exploits to support goal-directed behavior. In this framework, depth, motion, and somatosensory feedback contribute complementary channels of information that ultimately serve a common computational objective: transforming visual input into motor commands.

In summary, our model shows that central features of dorsal-ventral organization can arise from optimizing distinct behavioral objectives, even in the absence of recurrence, spatial topography, or motion inputs. Such constraints undoubtedly further sculpt the cortical architecture, but they need not be assumed for dual-stream structure to emerge. This supports the view that biological visual systems implement a general computational strategy, hierarchical feature refinement for perception and action, onto which evolution and development layer recurrence, anatomical constraints, and multimodal integration. Viewed this way, the dual-stream architecture is not a fixed design feature but a computational consequence of solving distinct goals in a structured world, one that biology subsequently elaborates and refines.

## Methods summary

Details of participants, dataset, model architecture, Shapley analyses, and EEG procedures are provided in the Online Methods.

## Online Methods

### Experimental Participants and Model Details

#### Human participants

Sixteen right-handed adults (age 21.56 ± 1.90; 37% male) with normal or corrected-to-normal vision participated in the study. All participants were students at the University of Toronto and provided written informed consent in accordance with the guidelines approved by the University of Toronto Research Ethics Board. Handedness was confirmed using a modified version of the Edinburgh Handedness Inventory^63^. Participants were compensated $120 for their time (8-9 hours total).

#### Dataset

We used a curated subset of the Jacquard dataset^32^ containing synthetic RGB-D images with detailed grasp annotations. From over 54,000 images, 1,000 were selected across five object classes (figurine, pen, chair, lamp, and plant; 250 per class). Only grasp annotations with confidence ≥ 0.8 were retained, yielding 30-40 valid candidates per image. All images were rendered on white backgrounds with objects shown in varied positions and orientations.

#### Neural Network Architecture

We implemented a multi-task convolutional neural network (Dual-task CNN) designed to jointly predict object class and grasp configuration from paired RGB-D inputs. The model takes as input 224 × 224 × 3 RGB and depth images, incorporating depth as an approximation of stereovision known to support human grasping^34^. The network comprises a feature extraction module and a feature expansion module (Fig. 1B).

In the feature extraction module, RGB and depth streams were each processed through the first convolutional block of a pretrained AlexNet^36^ (55 × 55 × 64), consisting of convolution, ReLU activation, and max pooling. These initial layers were frozen for the first 75 epochs to preserve pretrained low-level filters that capture early visual features analogous to V1 responses^36,64^. Outputs from both streams were concatenated (27 × 27 × 128) and passed through a convolutional block with a kernel size of 5, and stride of 1, (27 × 27 × 32), a max pooling layer with a kernel size of 3 and a stride of 2, then three additional convolutional layers (13 × 13 × 64) forming a shared visual representation that encodes mid-level features relevant for both grasping and object recognition.

The feature expansion module consisted of two parallel branches corresponding to grasping (dorsal) and classification (ventral) streams. Each branch employed convolution-deconvolution blocks to generate 224 × 224 × 6 pixel-wise output maps. The grasping stream predicted x-y coordinates of grasp centers, orientation (sin 2θ, cos 2θ), gripper width, and a confidence score. The classification stream produced category-specific activation maps (e.g., pen, lamp, plant, figurine, chair) plus a confidence channel, maintaining architectural symmetry with the grasping pathway. All layers used ReLU activations, transposed convolutions for spatial upsampling, and Xavier-uniform weight initialization^65^. The complete network contained 459,500 trainable parameters.

##### Loss Function

To enable consistent end-to-end training of both grasping and classification task under a unified architecture, we developed a novel single loss function called Double Log Loss (Figure 1C) The loss function preserves the spatial structure of the task while combining continuous and categorical objectives. This loss integrates a log-transformed L1 loss for regression targets and a negative log-likelihood (NLL) loss for confidence maps within one cohesive objective.

The total loss is computed as:

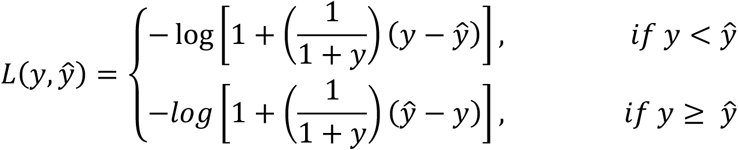

This formulation, used for the output channels, modulates error magnitude based on the ground truth value, effectively scaling the loss according to the proximity of predictions to their targets. It enhances stability over traditional L1 loss, particularly near boundary values (e.g., 0 or 1), and helps maintain informative gradients during backpropagation. By applying the logarithmic transformation, the loss further penalizes larger deviations while preventing vanishing gradients in near-correct predictions.

##### Model Training

The dataset was split into 90% training and 10% testing sets with object classes evenly distributed across both. The model was jointly trained on grasp prediction and object classification using paired RGB-D inputs, ensuring both outputs relied on a shared visual representation. Grasp supervision was provided through dense spatial maps encoding grasp centers, orientation, and gripper width, while classification supervision used category-specific maps and a confidence channel. Training used the Adam optimizer^66^ (learning rate = 5×10^-4^) for 150 epochs with a batch size of 5 in PyTorch^67^ 2.3.0 on NVIDIA RTX 1080 GPUs.

##### Evaluation Metrics

Model performance was assessed using task-specific criteria. For grasp planning, correctness followed the rectangle-based metric^68^: predictions were valid when the intersection-over-union (IoU) with the ground-truth grasp exceeded 25% and the orientation error was < 30°. This metric captures both spatial and directional accuracy, yielding 81% test accuracy on the Jacquard dataset. For object recognition, the element-wise products of the confidence map and each class activation map were calculated, enabling pixel-level classification while retaining spatial context. These products were then summed, with the class with the greatest sum being the model’s prediction. The model achieved 85% accuracy under this criterion.

#### Feature Map Correlations

To assess the contribution of individual feature maps to task performance, we applied the Neuron Shapley method, an approach derived from cooperative game theory and adapted for deep neural networks^39^. This method estimates the marginal contribution of each feature map (or “player”) to the model’s predictive accuracy by averaging its effect across multiple random subsets of the network, yielding a task-relevance score known as the Shapley value. Unlike activation- or gradient-based metrics, this approach directly quantifies how much each feature map contributes to overall model performance. This analysis was used to identify the feature maps most critical for grasping and classification, revealing how functional specialization emerged within a shared representational architecture.

For a given feature map *i*, its Shapley value *φ_i_* is defined as the expected marginal improvement in the model. Let *N* be the full set of feature maps, and *S* ⊆ *N*\{*i*} a subset excluding *i*. The Shapley value is defined as:

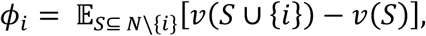

where *v*(*S*) denotes model’s performance using only feature map in *S*. This measures how much adding feature map *i* improves performance across different feature map combinations.

Because the number of subsets *S* grows exponentially with network size, we approximated this expectation using the Truncated Monte Carlo Shapley (TMC-Shapley^39,69^) algorithm. For each of *M* randomly sampled permutations *π* of the filter indices we computed the marginal contribution of feature map *i* as:

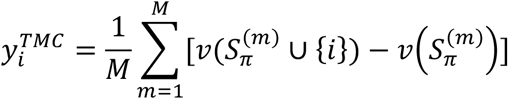

where 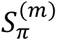 denotes the set of feature maps preceding *i* in the *mth* permutation. This stochastic approximation provides an unbiased estimator of the true Shapley value under reasonable assumptions about network modularity^39,69^.

To quantify the reliability of these Shapley estimates, we computed empirical confidence bounds using an empirical Bernstein inequality^70^, which accounts for both sample variance and the bounded range of the performance metric. For a given feature map *i*, the confidence bound *CB_i_* is given by:

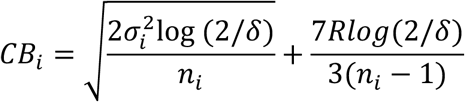

where 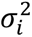 is the sample variance of contributions for feature map *i*, *n_i_* is the number of valid permutations, *R* is the bounded metric range, and *δ* = 0.05. Confidence intervals were visualized as error bars in layer-wise Shapley distributions to highlight high-impact, statistically robust units. Shapley values were then used to rank feature maps by task relevance.

To assess how task relevance co-varied between grasping and classification, Pearson correlations between Shapley values of the two tasks were computed for each shared convolutional layer. Correlations were bootstrapped over 1,000 resamples to obtain mean *r*-values and 95% confidence intervals^71^. These correlation profiles captured the degree of functional overlap across tasks and revealed decreasing cross-task correspondence with network depth, consistent with increasing functional specialization.

#### Activation Maximization

To visualize the preferred stimulus features of the most task-relevant units, we used activation maximization, an optimization technique that iteratively adjusts an input image to maximally activate a given feature map^72^. This approach produces synthetic patterns that reveal the visual features each unit is tuned to, analogous to mapping neuronal receptive fields. The resulting images highlighted distinct feature preferences for grasp-and classification-selective maps, linking model representations to cortical feature selectivity.

#### Weight Analysis

To complement the feature-map level Neuron Shapley analysis^38^, we applied the Shapley Value Approximation without Marginal contributions (SVARM) algorithm^45^ to estimate task relevance at the weight level. Whereas Neuron Shapley samples permutations of feature maps and becomes computationally prohibitive for large parameter sets, SVARM approximates Shapley values analytically through coalition sampling, allowing tractable estimation across millions of weights while revealing fine-grained contribution structure within each layer.

We randomly sampled 250 test images and ran SVARM twice - once for classification and once for grasp prediction - across all five shared layers. In each run, the players were the weights in that layer. For a weight *i*, its contribution is computed as the difference between two weighted averages of coalition performances - with and without that weight:

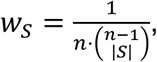

where *n* is the number of weights in the layer and *S* a coalition of retained weights.

Coalitions are sampled from:

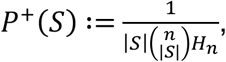

and

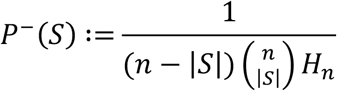

Here, *H_n_* is the n^th^ harmonic number. When a coalition *A*^+^ is sampled from *P*^+^(*S*), the positive Shapley values for all players *i* ∈ *A*^+^ are updated, and likewise when a coalition *A*^−^ is sampled from *P*^−^(*S*), the negative Shapley values for all players *i* ∉ *A*^−^ are updated. Theoretical results for SVARM show that the expected mean squared error between estimated Shapley value and true Shapley value is bounded by,

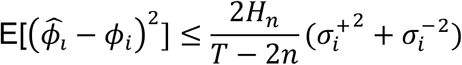

where *T* is the total number of iterations, and 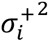 and 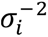 are the variances of player *i*’s positive and negative Shapley value, respectively. We used this bound in conjunction with the variance of the Shapley values across all players to inform the choice of *T* that would suffice for a meaningful analysis. In practice, we ran the SVARM algorithm for each layer and task combination until the fraction of variance in Shapley values due to estimator noise was ≤ 0.1-0.2.

We computed Pearson correlations between classification and grasping Shapley values across kernels for each layer. Confidence intervals for the true correlation were estimated using theoretical MSE bounds combined with nonparametric bootstrap resampling. For each task, the reliability lower bound was defined as 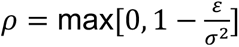 where *ε* is the average expected MSE bound and σ^2^ the sample variance. The true correlation interval was obtained via classical disattenuation^73^ and refined through 1,000 bootstrap resamples, with the 95% bounds given by the 2.5th and 97.5th percentiles of the bootstrapped estimates.

#### Double Dissociation Analysis

To test whether the shared layers contained functionally specialized representations for grasping and classification, we conducted a double dissociation analysis based on systematic feature map lesioning. Within each shared convolutional layer, feature maps were ranked by task relevance using Neuron Shapley values and progressively silenced. Two lesion conditions were tested: *grasp-impact lesions*, in which grasp-selective feature maps were set to zero, and *classification-impact lesions*, in which classification-selective maps were set to zero. We limited lesions to 50% of feature maps within a layer to avoid producing non-specific deterioration throughout the network. For each lesion and layer, model performance was evaluated across 250 randomly sampled test images, yielding distributions of accuracy for both tasks at each lesion proportion.

To quantify selective impairment, we computed *t*-values comparing post-lesion accuracy to the intact baseline for each task and lesion type. A double dissociation was identified when lesioning feature maps specialized for one task significantly reduced performance in that task (*t* > 2.33, *p* < 0.01) while sparing the other. This analysis revealed layer-specific patterns of selective disruption, demonstrating that different subnetworks contributed independently to grasping and object classification within the shared architecture.

#### Functional Connectivity Analysis

To examine how task-selective feature maps interacted across layers, we modeled network connectivity using directed, weighted graphs. For each pair of adjacent convolutional layers, Pearson correlations between flattened kernel weight vectors were computed to form inter-layer adjacency matrices capturing connection strength between feature maps. Each node (feature map) was categorized as grasp-selective, classification-selective, or no preference based on normalized Neuron Shapley values, where maps exceeding the 80th percentile of task-specific Shapley distributions were assigned to their respective task and all others were labeled as no preference. Edge weights were normalized within each layer pair and thresholded at 1.2× the mean inter-layer weight to retain the most informative connections. Within-task links were shown as solid red or blue edges, while purple dashed edges indicated cross-task connections between grasp- and classification-selective nodes.

#### Connection Analysis and Visualization

Using feature-map-level Shapley values, we examined how task-specific connections between layers evolved across the network. Feature maps were first classified as classification-specialized, grasp-specialized, or nonselective based on their relative task relevance using two complementary metrics: the effect-size measure (*d^* = 0.5 or 0.15) and the top-k criterion (k = 20 and k = 30). For each pair of adjacent layers, we identified all possible connections between feature maps and labeled each connection according to the source and target map types - classification (both classification-specialized), grasp (both grasp-specialized), cross (one of each), or nonselective (involving at least one neutral map).

The pie charts in Fig. 4B show the proportion of these four connection types for each layer pair, illustrating how functional specialization shifts as the network deepens. To statistically test whether deeper layers contained a higher fraction of specialized or cross-task connections, we created weighted contingency tables comparing specialized versus non-specialized connections across consecutive layer pairs and conducted weighted χ² tests. Connection counts were weighted by the magnitude of the mean of the connection’s kernel. The resulting χ² values (Fig. 4C) quantify the magnitude and significance of these changes for both the effect-size and top-k metrics.

#### Representational Similarity Analysis (CNN-EEG Correlation)

We used representational similarity (or dissimilarity) analysis (RSA or RDA)^74^ to examine how the internal representations of a single convolutional neural network (CNN) trained on grasping and object classification aligned with time-resolved EEG activity recorded during task performance. During both model inference and EEG recording, identical images of 25 previously unseen objects were presented, ensuring out-of-sample evaluation of representational geometry.

##### Extraction of CNN Representations

For the Dual-Task-CNN, we extracted activation patterns from the first five convolutional layers during inference. A forward-pass hook-based method was used to record the output tensors of the selected layers without modifying network architecture or weights. Formally, let 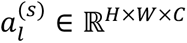 denote the activation tensor of layer *l* for stimulus *s*, where *H*, *W* and *C* are the spatial height, width, and number of channels, respectively. These activations were flattened into feature vectors 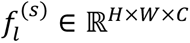 and z-scored across stimuli. A representational dissimilarity matrix (RDM) was then computed for each layer using pairwise correlation distances across all object pairs:

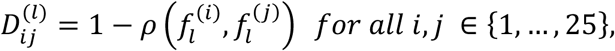

where *ρ*(.,.) is Pearson’s correlation coefficient. This procedure yielded a single 25×25 RDM per layer for each model, capturing layer-specific representational geometry across stimuli.

##### EEG-Based Representational Geometry

EEG signals were recorded while participants performed either a grasping or classification task with the same 25 stimuli. Trials were epoched relative to stimulus onset and segmented into contiguous, non-overlapping 25 ms windows from -25 ms to 400 ms. Within each time window *t*, the evoked response 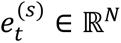 was computed by averaging across trials and across *N* posterocentral electrodes (CP, P, PO and O) for each stimulus *s*. These electrodes were selected because they overlie visual and posterior parietal cortices and show the strongest early visuomotor and shape-related responses, making them optimal for capturing object-driven dynamics relevant to both tasks. To remove baseline bias, we then subtracted the grand mean across stimuli from each response vector:

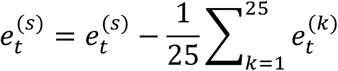

As with the model activations, we next computed an RDM for each time window using correlation distance:

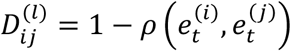

This yielded one EEG RDM per time window, capturing the evolving representational geometry of the cortical response.

##### Model-to-EEG Correlation

To quantify the alignment between CNN layer *l* and EEG time bin *t*, we computed the Spearman’s rank correlation coefficient between the upper triangular portions of each model RDM *D*^(*l*)^ and each EEG RDM *D*^(*t*)^:

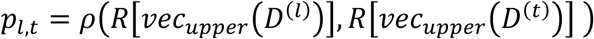

where R[*x*] converts raw score *x* to its corresponding rank variable. Thus, *p_l,t_* denotes the similarity between layer *l* of the CNN and EEG responses at time window *t*. This produced a layer-by-time matrix **R** ∈ ℝ^5×*t*^ of correlation values for each model (grasp and classification), where T represents the number of time bin.

To assess the reliability of each correspondence value *p_l,t_*, we conducted a non-parametric permutation test^75^ in which EEG RDM stimulus labels were randomly shuffled 10,000 times for each time window to generate a null distribution. Correlations exceeding the 95th percentile of this distribution were deemed significant. The resulting layer-by-time correlation matrices were visualized as bar plots, enabling inspection of how the representational geometry of the CNN aligned with cortical dynamics across time and network depth. This analysis provided a fine-grained temporal mapping between CNN-derived representations and neural activity, revealing how different stages of model processing corresponded to evolving brain responses during grasping and object recognition^54,76^.

## Competing interests

The authors declare no competing interests.

## Funding

This work was supported by NSERC Discovery Grant RGPIN-2020-06018 (to M. Niemeier)

## Notes

### Competing Interest Statement

The authors have declared no competing interest.

### Summary of Updates

Details added to the Methods section

